# Glutamatergic synaptic inhibition through group II mGluR-mediated suppression of the sodium leak channel NALCN

**DOI:** 10.64898/2026.08.21.746377

**Authors:** Christian T. Candler, Kennedy E. Whittaker, Timothy S. Balmer

## Abstract

The sodium leak channel NALCN regulates resting membrane potential and spontaneous firing in neurons and can be modulated by G-protein coupled receptors (GPCRs). Whether metabotropic glutamate receptors (mGluRs) modulate NALCN is unknown and would represent a novel mechanism through which glutamate could affect neuronal excitability. Here we examine NALCN function and modulation by mGluRs in cerebellar unipolar brush cells (UBCs) in mouse brain slices. Activation of group II mGluRs inhibited the NALCN current through a G protein-dependent mechanism, as the effect was abolished by intracellular GDP-β-S and by NALCN deletion. The OFF UBC subtype that is inhibited by glutamate had a larger NALCN current than the ON UBC subtype that is excited by glutamate. OFF UBCs also had a tonic NALCN current that was absent in ON UBCs. Genetic deletion of NALCN converted the regular spontaneous firing pattern of OFF UBCs, to an irregular pattern similar to that of ON UBCs, suggesting that a tonic NALCN current may be a general mechanism to promote regular firing. Additionally, we identify the presence of group III mGluRs in OFF UBCs and GABA-B receptors in ON UBCs and show that neither inhibit NALCN, demonstrating that different GPCRs engage distinct downstream ion channels. These findings identify a previously unrecognized form of glutamatergic synaptic inhibition that is selectively initiated by group II mGluRs, but not other Gi/o-coupled GPCRs, within the same neurons.

**Graphical Abstract:** 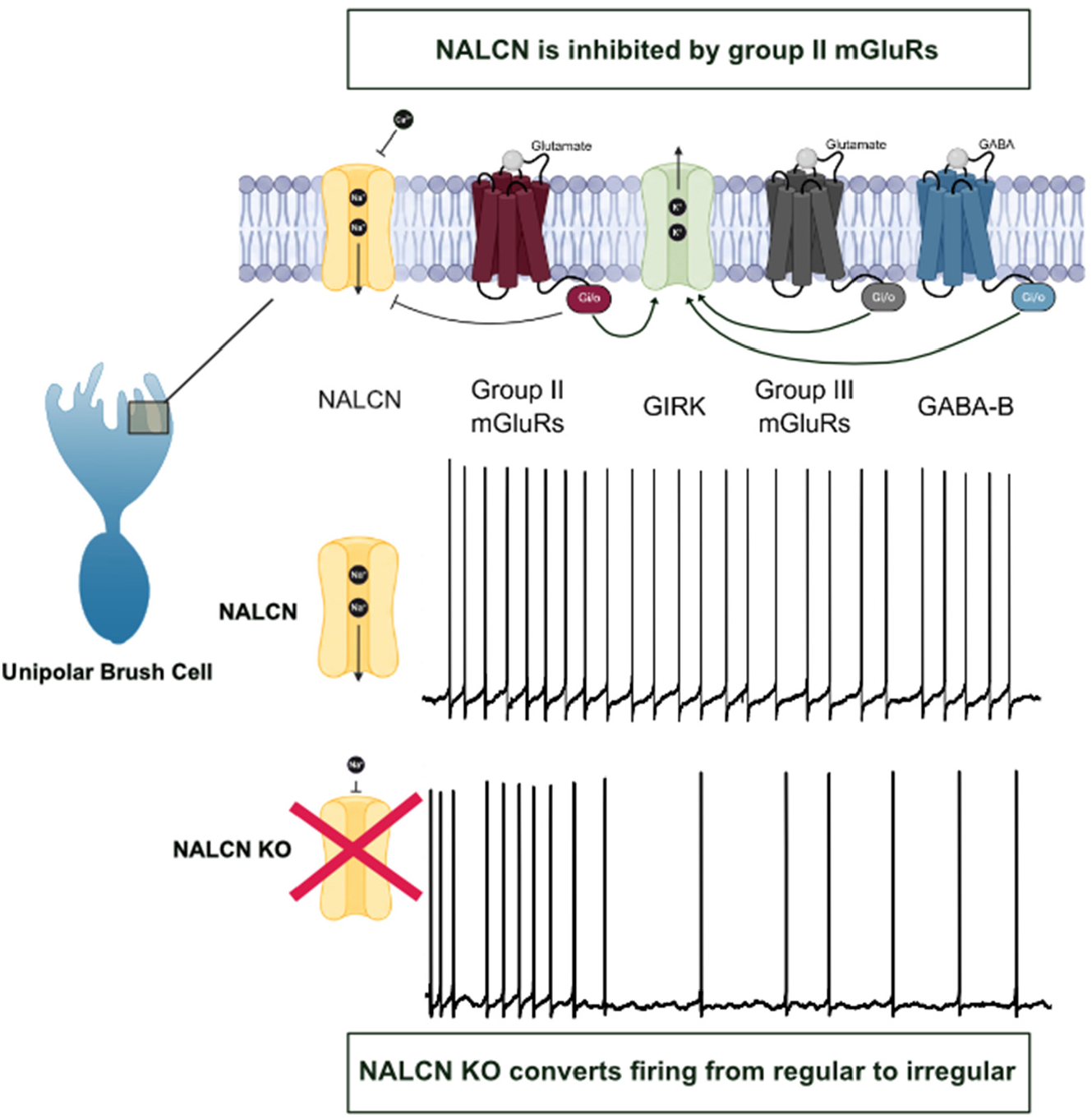

## Introduction

Metabotropic neurotransmitter receptors release intracellular G-proteins that interact with various ion channels to regulate cellular excitability (Dascal, 2001). One such ion channel, the Na^+^ leak channel NALCN (sodium leak channel nonspecific), controls excitability by depolarizing the resting membrane potential (Lu *et al*., 2007; Lutas *et al*., 2016).

NALCN has been shown to regulate pacemaking (Lu *et al*., 2007; Um *et al*., 2021), motor activity (Flourakis *et al*., 2015), and circadian rhythms (Gonzalez *et al*., 2023), and its dysfunction can contribute to developmental disorders including cerebellar ataxia in humans (Aoyagi *et al*., 2015; Kschonsak *et al*., 2022; Cabrita Pinto *et al*., 2025). The conductance of NALCN can be regulated by multiple intracellular signaling pathways (Philippart & Khaliq, 2018; Hahn *et al*., 2020; Um *et al*., 2021). For example, GABA-B and dopamine receptors release G-proteins (Gi/o) that inhibit NALCN, in addition to activating inwardly rectifying potassium channels (GIRK) (Philippart & Khaliq, 2018; Cobb-Lewis *et al*., 2023; Ngodup *et al*., 2024). It is clear that group II and group III metabotropic glutamate receptors (mGluR2/3 and mGluR4/6/7/8, respectively) hyperpolarize neurons by releasing Gi/o that activates GIRK (Kammermeier *et al*., 2003; Niswender *et al*., 2008). However, it remains unknown whether these inhibitory mGluRs also modulate NALCN currents. If this is the case, it would represent a novel mechanism through which glutamate could produce synaptic inhibition.

Here we address the role of NALCN in intrinsic excitability and synaptic inhibition using unipolar brush cells (UBCs), which are glutamatergic interneurons in cerebellum-like circuits (Mugnaini *et al*., 2011). RNA-sequencing has shown that UBCs express NALCN channel complex transcripts as well as multiple types of Gi/o-coupled receptors that could modulate NALCN activity (Saunders *et al*., 2018; Kozareva *et al*., 2021; Jing *et al*., 2025). UBCs can be divided into two subtypes based on their response to glutamate and their spontaneous firing patterns: ON UBCs are excited by excitatory mGluR1 and AMPA receptors and fire irregularly, and OFF UBCs are inhibited by group II mGluRs and fire regularly (Russo *et al*., 2007; Kim *et al*., 2012; Borges-Merjane & Trussell, 2015; Balmer *et al*., 2021). Because NALCN can promote rhythmic firing (Lu *et al*., 2007; Lutas *et al*., 2016; Hahn *et al*., 2023; Cobb-Lewis *et al*., 2023; Ngodup *et al*., 2024), we hypothesized that its differential expression across the UBC population could relate to their different spontaneous firing patterns. Tonic depolarization mediated by NALCN may represent a general mechanism for promoting regular spontaneous firing, thereby priming neurons for specialized roles in sensory processing that distinguish them from neurons with irregular firing patterns. (Goldberg & Fernández, 1977; Goldberg *et al*., 1984; Eatock *et al*., 2008).

The properties of UBCs allowed us to test (1) how NALCN contributes to spontaneous firing patterns, (2) the role of NALCN in mediating synaptic inhibition by glutamate, and (3) whether different Gi/o-coupled GPCRs target different channels in the same cells. We show that NALCN is essential for setting cellular excitability and driving regular spontaneous firing patterns, which may be essential for the different roles of ON and OFF UBCs in signal processing in cerebellum-like circuits. Glutamate can hyperpolarize neurons through group II mGluR-mediated inhibition of a tonic NALCN current, in addition to its role in activating GIRK. Group III mGluRs and GABA-B receptors act through GIRK but do not inhibit NALCN, indicating the presence of a distinct intracellular pathway for group II mGluRs. mGluR-mediated inhibition of NALCN represents a newly identified form of synaptic inhibition by glutamate that may occur across the nervous system.

## Methods

### Animals

Adult mice of both sexes at postnatal age (P22-82) were used. Different mouse lines were generated to label ON and OFF UBCs specifically. The P079 line: Et(tTA/mCitrine)P079Sbn (Shima *et al*., 2016; Hariani *et al*., 2024) labeled OFF UBCs, and the GRP-Cre line: Tg(Grp-Cre)KH107Gsat (MMRRC_031182-UCD) (Kim *et al*., 2012; Gerfen *et al*., 2013; Hariani *et al*., 2024) crossed with a tdTomato reporter: Ai9 Gt(ROSA)26Sor^tm9(CAG-tdTomato)Hze^ (IMSR_JAX:007909)(Madisen *et al*., 2010) labeled ON UBCs. A NALCN conditional knockout line: B6(Cg)-Nalcn^tm1c(KOMP)Wtsi/DrenJ^ (IMSR_JAX:030718) (Yeh *et al*., 2017) was crossed with a tamoxifen-inducible calretinin-Cre line: Calb2^tm2.1(cre/ERT2)Zjh/J^ (IMSR_JAX:013730) (Taniguchi *et al*., 2011) and with a tdTomato reporter line: Ai9 Gt(ROSA)26Sor^tm9(CAG-tdTomato)Hze^ (IMSR_JAX:007909) (Madisen *et al*., 2010). Cre-mediated recombination was induced in calretinin-Cre mice by intraperitoneal injections of 10 mg/ml tamoxifen (T5648, Sigma) in corn oil at a dose of 75 mg/kg per day for 3 days and electrophysiological experiments were performed 2 weeks later. Mice were bred in a colony maintained in the animal facility managed by the Department of Animal Care and Technologies, and all procedures were approved by Arizona State University’s Institutional Animal Care and Use Committee under protocol #24-2044R. Transgenic mice were genotyped by light at P0– P3, or toes were clipped at P9 and genotyped by PCR.

### Acute brain slice preparation

Mice were deeply anesthetized with isoflurane which was confirmed with a toe pinch. The brain was extracted and glued to a platform on a vibratome (Campden Instruments 7000smz-2) and cut in 200 or 300 μm thick sagittal sections in ice-cold cutting solution containing (in mM) 87 NaCl, 2.5 KCl, 1.25 NaH_2_PO_4_, 0.4 Na-Ascorbate, 2 Na-Pyruvate, 25 Glucose, 25 NaHCO_3_, 75 Sucrose, 7 MgCl_2_, 0.5 CaCl_2_. Slices were then transferred to ACSF at 35°C for 35 minutes before maintaining at room temperature (∼23°C) until recording. Cutting solution and ACSF were bubbled with 95% O_2_ / 5% CO_2_. Recordings were made from Lobes IX and X of the cerebellum within 8 hours of dissection.

### Artificial cerebral spinal fluid (ACSF)

Control ACSF included (in mM) 130 NaCl, 2.1 KCl, 1.2 KH_2_PO_4_, 0.4 Na-Ascorbate, 2 Na-Pyruvate, 3 Na-HEPES, 1 MgSO_4_ 10 Glucose, 20 NaHCO_3_, and 0.1 or 1.5 CaCl_2_. Osmolarity was between 300-310 with pH ∼7.3. To isolate NALCN currents and block GIRK, the following drugs were added to the ACSF: 3 mM Cs^2+^, 0.5 µM TTX, 0.3 µM Tertiapin-Q, 100 μM Ba^2+^, 1 μM JNJ16259685. Other drugs were bath applied as noted in the results section. The NMDG solution was made by dissolving (in mM) 130 NMDG, 2.1 KCl, 1.2 KH_2_PO_4_, 0.4 Na-Ascorbate, 2 Na-Pyruvate, and 25 NaHCO_3_ in ddH_2_0, adjusting the pH to 7.3-7.4 with HCl, and then adding 10 Glucose, 1 MgSO_4,_ and 0.1 or 1.5 CaCl_2_.

### Internal pipette solutions

For voltage clamp experiments to isolate the NALCN current, the internal pipette solution contained (in mM) 145 CH_3_O_3_SCs, 10 QX-314, 2 MgCl_2_, 5 K_2_ATP, 0.5 EGTA, 5 HEPES; adjusted to 290 mOsm with sucrose, and pH-ed with CsOH to 7.2-7.25. For current clamp experiments and all other voltage clamp experiments, the internal solution contained (in mM) 113 K-gluconate, 9 HEPES, 4.5 MgCl2, 0.1 EGTA, 14 Tris-phosphocreatine, 4 Na_2_-ATP, 0.3 Tris-GFP, with 0.3% biocytin. pH was adjusted to 7.2-7.3 with KOH, and osmolarity was adjusted to 290 mOsm with sucrose. Reported voltages were corrected for 15-mV and 12-mV liquid junction potentials for the Cs-methylsulfonate and K-gluconate internal solutions, respectively.

### Acute brain slice electrophysiology

Slices were recorded on a fixed-stage microscope (Olympus BX51) with Dodt gradient contrast optics (Scientifica), and a 60X water immersion Olympus objective. 6-10 MΩ patch pipettes were pulled from borosilicate glass capillaries (OD 1.2 mm and ID 0.68 mm, AM Systems or Sutter) with a horizontal puller (P1000, Sutter Instruments). Data were acquired using a Multiclamp 700B amplifier and pClamp 11 software (Molecular Devices). Signals were acquired with 5-10X gain, sampled at 50-100 kHz using a Digidata (1550A, Molecular Devices) analog-digital converter, and low-pass filtered at 10 kHz. Series resistance was compensated with correction 20-40% and prediction 50-70%, bandwidth 2 kHz. Additional digital filtering was applied o?ine (Bessel-1 kHz for noise analysis).

### Data Analysis

Clampfit (Molecular Devices) and Prism (Graphpad) were used for analysis. Most statistical significance in this study was determined by paired or unpaired t-tests with Welch’s correction. All data are reported in mean ± SEM. For the firing pattern analysis, the interspike intervals during 60-second recordings from each cell were put into 10-ms bins to produce histograms. A Kolmogorov-Smirnov test was used to compare interspike intervals between the wild-type and NALCN knock-out mice.

## Results

### OFF UBCs have a larger NALCN current than ON UBCs

RNA sequencing previously showed high expression levels of NALCN channel complex genes (*Nalcn, Fam155a, Unc79, Unc80*) in UBCs, but the analysis did not differentiate between ON and OFF UBCs (Saunders *et al*., 2018; Kozareva *et al*., 2021; Jing *et al*., 2025). To test whether the NALCN current differed between ON and OFF UBCs, we ran a series of whole cell patch clamp electrophysiology experiments in both subtypes, defined by their labeling in transgenic mouse lines: P079-previously reported to label OFF UBCs, and GRP-Ai9-previously reported to label ON UBCs (Gerfen *et al*., 2013; Shima *et al*., 2016; Hariani *et al*., 2024). To isolate NALCN currents, TTX was used to block a Na^+^ current through voltage gated Na^+^ channels that is present in some UBCs (Balmer *et al*., 2021). NALCN is blocked by extracellular Ca^2+^ (Lu *et al*., 2010; Chua *et al*., 2020), so reducing the Ca^2+^ concentration in the ACSF produces an inward current in voltage-clamp that can be used as a measure of the total NALCN current in the cell. Extracellular Ca^2+^ was lowered from 1.5 to 0.1 mM and an inward current was observed in both ON (-34.12 ± 5.364 pA, n = 10, paired t-test, p = 0.0001)) and OFF UBCs (-96.6 ± 12.04 pA, n = 13, p = 0.0013) (**Fig. 1A-B**). Reducing the extracellular Na^+^ concentration from 144 mM to 14 mM by replacing it with a solution containing NMDG confirmed that Na^+^ was the charge carrying ion in both ON (16.47 ± 3.398 pA, n = 10, paired t-test, p = 0.0009) and OFF UBCs (122.9 ± 46.26 pA, n = 7, paired t-test p = 0.0039). The NALCN current in OFF UBCs was significantly larger than the NALCN current in ON UBCs (unpaired t-test with Welch’s correction, p < 0.001, **Fig. 1C**).

**Figure 1:**
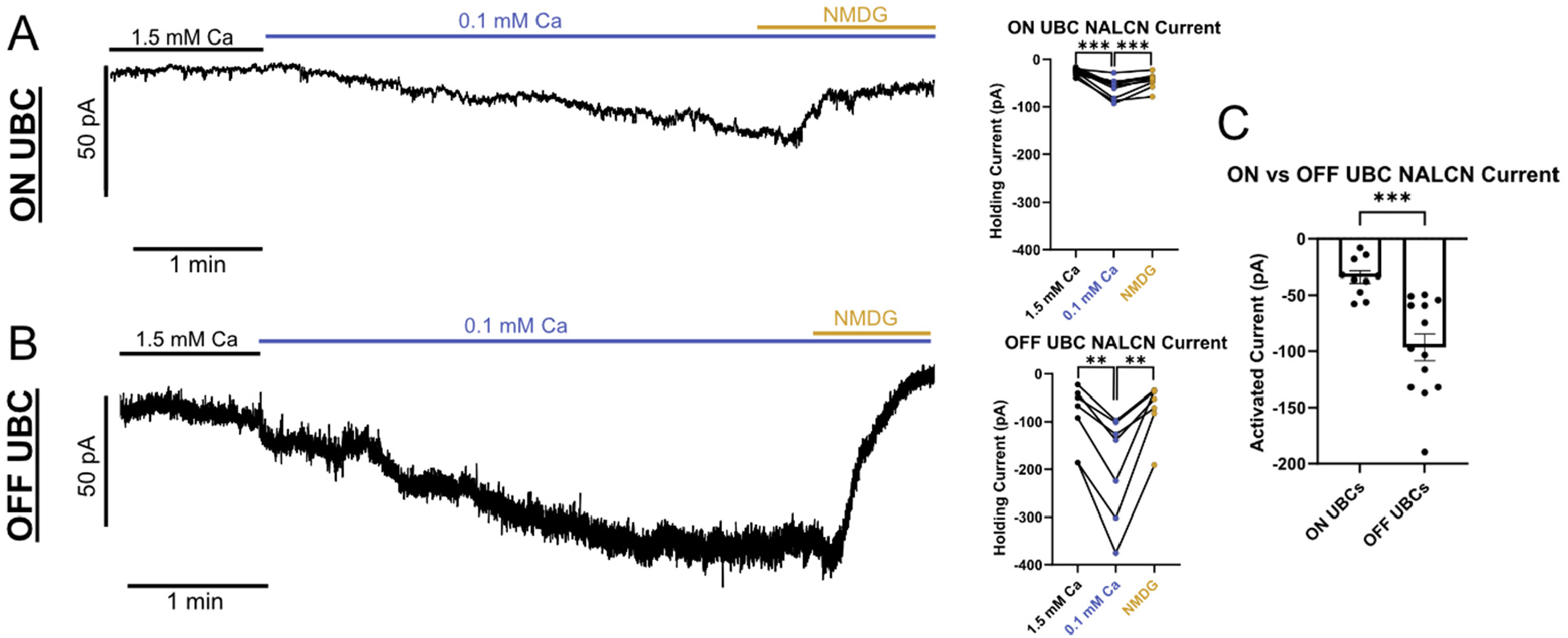
OFF UBCs have a larger total NALCN current than ON UBCs. A) Left-voltage clamp recording of NALCN current in an ON UBC. The Ca^2+^ concentration of the bath solution was lowered to 0.1 mM and a depolarizing inward current was observed. Sodium was replaced with NMDG to remove the charge carrier, which reduced the inward current, shifting it outward. Right-summary data. B) Left-voltage clamp recording of NALCN current in an OFF UBC with solution exchanges as in A. Right-summary data. C) The NALCN current was significantly larger in OFF UBCs than in ON UBCs.

Many neurons are tonically depolarized by an active NALCN conductance at rest. To test whether UBCs have a tonic NALCN current, we recorded from UBCs in 1.5 mM Ca^2+^ and reduced the extracellular Na^+^ concentration with an NMDG solution. ON UBCs showed no significant change in holding current (3.046 ± 2.319 pA, n = 10, paired t-test, p = 0.2216, **Fig. 2A**), while OFF UBCs showed an outward shift in the holding current (22.2 ± 5.525 pA, n = 7, paired t-test, p = 0.007, **Fig. 2B**). In a few ON UBCs there was a modest outward shift in the holding current indicating some may have a small tonic NALCN current. However, OFF UBCs had a significantly larger tonic NALCN current than ON UBCs (unpaired t-test with Welch’s correction, p = 0.034, **Fig. 2C**). Taken together, these data show that OFF UBCs have a large NALCN current that produces a significant depolarizing tonic current and ON UBCs have a smaller NALCN current that produces little to no tonic current.

**Figure 2:**
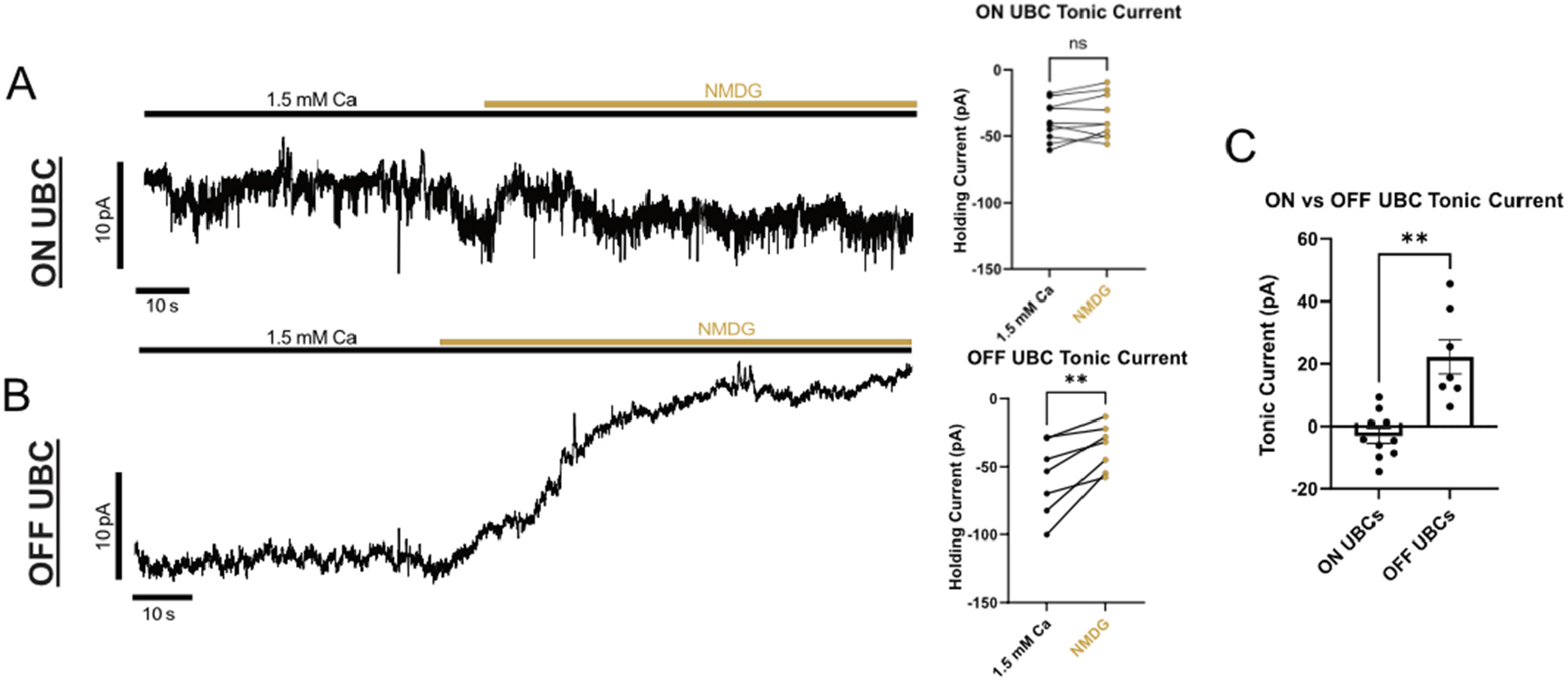
OFF UBCs have a larger tonic current than ON UBCs. A) Left-ON UBC tonic NALCN current was measured in voltage clamp by replacing 1.5 mM Ca^2+^ ACSF with an NMDG solution. Right-summary data. B) Left-OFF UBCs tonic NALCN current was measured in voltage clamp by replacing 1.5 mM Ca^2+^ ACSF with an NMDG solution. This produced an outward current, indicating a tonic inward NALCN current. Right-summary data. C) The tonic NALCN current was significantly larger in OFF UBCs than in ON UBCs.

### NALCN regulates pacemaking in OFF UBCs

NALCN regulates rhythmic activity and pacemaking across neurons and cardiac cells (Lu *et al*., 2007; Flourakis *et al*., 2015; Lutas *et al*., 2016; Shi *et al*., 2016; Philippart & Khaliq, 2018; Um *et al*., 2021; Cobb-Lewis *et al*., 2023). To test if NALCN regulates pacemaking in OFF UBCs, we generated a conditional NALCN knockout (KO) specifically in calretinin-expressing OFF UBCs (Calb2-CreERT2/Ai9/NALCN^flox/flox^). 2-3 weeks after tamoxifen injections we compared the spontaneous firing of wild-type (WT) and NALCN KO OFF UBCs. More hyperpolarizing current was required to hold the cell at -72 mV in voltage clamp in WT OFF UBCs (-54.34 ± 6.211 pA, n = 18) than in NALCN KO OFF UBCs (-27.02 ± 2.301 pA, n = 12, unpaired t-test, p = 0.0005, **Fig. 3A**), consistent with a tonic depolarizing current mediated by NALCN in WTs that was not present in KOs. The WT OFF UBCs fired significantly more (n = 5652 total spikes, 6 cells, 16.09 ± 5.189 Hz) than NALCN KO UBCs (n = 825 total spikes, 6 cells, 1.863 ± 1.140 Hz, Mann-Whitney non-parametric t-test, p = 0.0152), owing to the lack of tonic depolarizing current in the absence of NALCN (**Fig. 3B**). Three out of six NALCN KO UBCs were so hyperpolarized that they did not fire during the 60 second recordings.

**Figure 3:**
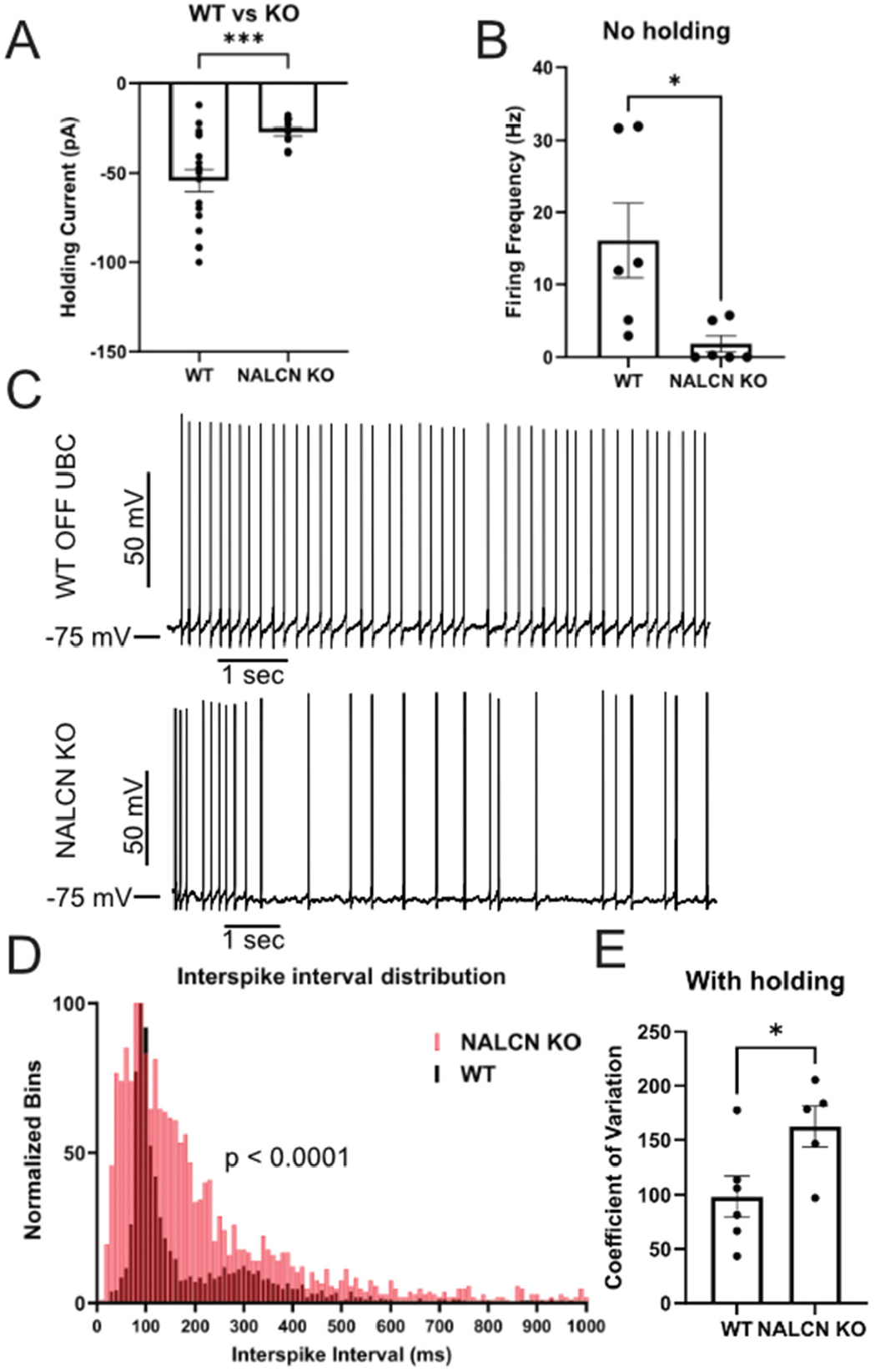
NALCN regulates spontaneous firing in OFF UBCs. A) Upon break-in, NALCN KO OFF UBCs required less negative current to hold the cell at -72 mV than WT UBCs. B) The firing frequency of NALCN KO OFF UBCs was much lower than WT UBCs. C) Representative traces from WT OFF UBC and NALCN KO OFF UBC in current clamp. Holding current was applied to maintain the cells at a resting membrane potential around -75 mV. D) Interspike interval histogram of spontaneous spikes in WT (black) and NALCN KO (red) OFF UBCs in 10-ms bins. E) The interspike interval of WT UBCs was less variable than that of NALCN KO UBCs.

When a bias current was applied to maintain the resting membrane potential around -75 mV, WT OFF UBCs had less variability in their interspike intervals compared to NALCN KO OFF UBCs (Kolmogorov-Smirnov test, p < 0.0001, **Fig. 3C-D**). WT OFF UBCs (3.671 ± 1.372 Hz, n = 6 cells) and NALCN KO UBCs (2.969 ± 0.8701 Hz, n = 6 cells0 had similar firing frequencies when they were held around -75 mV (unpaired t-test, p = 0.9307). WT UBCs had a significantly lower coefficient of variation (98.28 ± 19.02, n = 6 cells) compared to NALCN KO OFF UBCs (162.4 ± 18.81, n = 6 cells, unpaired t-test, p = 0.0404, **Fig. 3E**). These data indicate that the tonic NALCN current depolarizes OFF UBCs and may promote their spontaneous firing pattern.

### Glutamate-evoked outward current is mediated by GIRK and NALCN in OFF UBCs

Previous work has shown that inhibitory G-proteins released by activation of D2 dopamine receptors and GABA-B receptors can inhibit NALCN (Philippart & Khaliq, 2018; Gonzalez *et al*., 2023; Ngodup *et al*., 2024). Group II mGluRs also activate intracellular inhibitory G-proteins (Kammermeier *et al*., 2003). We hypothesized that activating group II mGluRs would release G-proteins that would inhibit NALCN in UBCs, along with their known intracellular target, GIRK.

We puffed glutamate onto OFF UBCs to evoke an inhibitory outward current in the presence of 10 µM NBQX and 1 µM JNJ16259685 to block AMPA/kainate and mGluR1 to isolate the group II mGluR current (**Fig. 4A**). 100 µM Ba^2+^ was applied to partially block GIRK (paired t-test, p < 0.0001, n = 8), followed by 10 µM L-703,606, a NALCN channel blocker, which also significantly decreased the current amplitude (paired t-test, p = 0.0155, n = 6). This result suggests that glutamate can have an inhibitory action by suppressing a tonic inward NALCN current.

**Figure 4:**
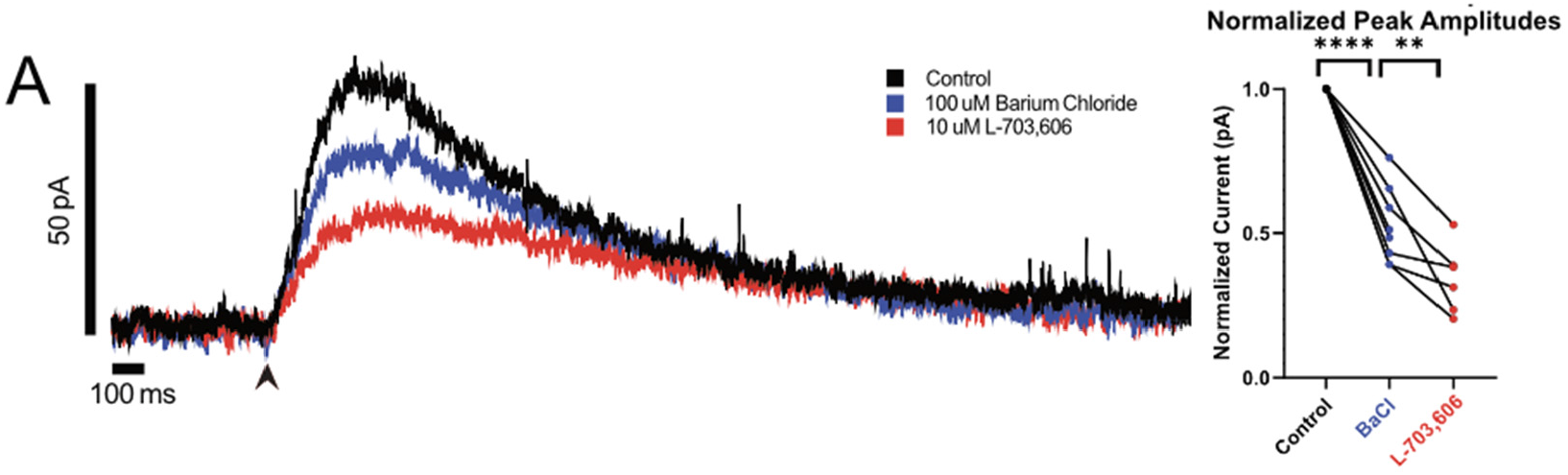
Glutamate evoked outward current is mediated by GIRK and NALCN in OFF UBCs. A) Left-A 10-ms glutamate (1 mM) puff induced an outward current in an OFF UBC. The amplitude of the current was reduced by bath application of Ba^2+^ to block GIRK,, and was further reduced by the NALCN channel blocker. Right-summary data.

Next, we tested whether the NALCN current is inhibited by group II mGluRs. A large NALCN current was induced by reduction of extracellular Ca^2+^ in the presence of GIRK blockers (3 mM Cs^2+^, 0.3 µM Tertiapin-Q, 100 μM Ba^2+^, intracellular Cs^2+^) and the group II mGluR agonist LY354740 was applied to test whether it reduced the NALCN current **(Fig. 5A)**. An NALCN current was activated by reducing the extracellular Ca^2+^ concentration (-56.45 ± 14.31 pA, n = 11), which was significantly reduced by 1 µM LY354740 (43.33 ± 10.94 pA, n = 11, paired t-test, p = 0.027, **Fig. 5A**).

**Figure 5:**
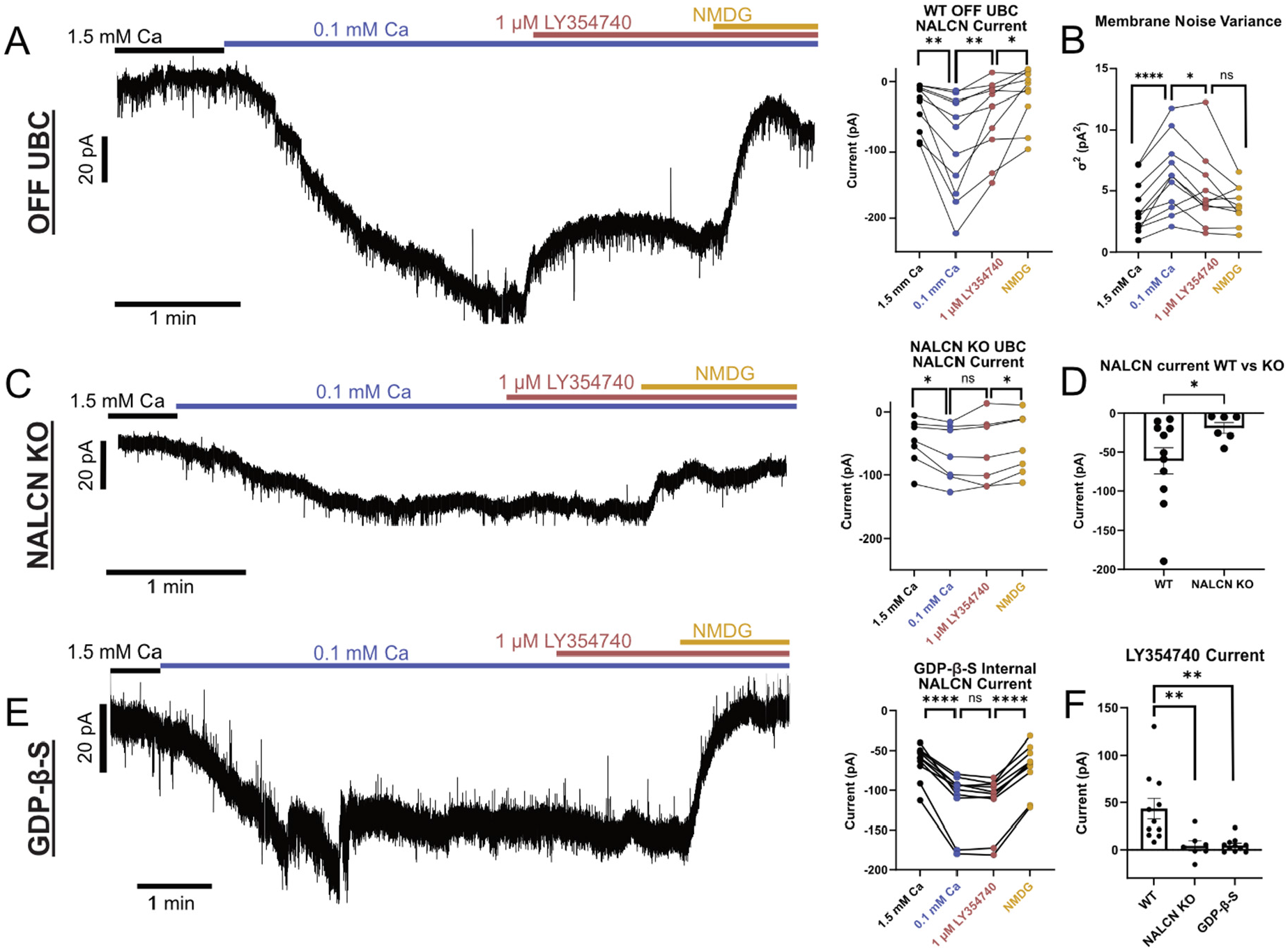
Group II mGluR activation inhibits NALCN current. A) Left-voltage clamp recording of an OFF UBC. The extracellular Ca^2+^ concentration was lowered to 0.1 mM to activate NALCN, then 1 μM LY354740 (group II mGluR agonist) was applied to the bath, followed by the NMDG solution. Right-Summary data. The group II mGLuR agonist significantly inhibited the low calcium activated inward NALCN current in the presence of GIRK blockers. B) Comparison of the membrane noise variance across each condition. Noise variance increased significantly in 0.1 mM Ca^2+^ and was suppressed by application of LY354740. C) Left-same experiment as above but in an NALCN KO OFF UBC. Right-lowering Ca^2+^ produced a small inward current and LY354740 did not have a significant effect on UBCs lacking NALCN. D) NALCN KO UBCs had a significantly smaller low Ca^2+^ induced NALCN current than wild type OFF UBCs. E) Left-same experiment as in A and B, but with addition of 10 mM GDP-β-S in the internal pipette solution. LY354740 did not have a significant effect on the NALCN current. Right-summary data. F) Comparison of the change in current amplitude after application of LY354740 in the conditions in A-C. Current amplitude in WT UBCs was significantly different compared to the NALCN KO UBCs or when GDP-β-S was in the internal pipette solution.

Transitioning into the NMDG solution to reduce extracellular Na^+^ reduced the NALCN current further (30.71± 11.35 pA, n = 11, paired t-test, p = 0.0242, **Fig. 5A**). Membrane noise variance (σ^2^) increased in 0.1 mM Ca^2+^ compared to baseline 1.5 mM Ca^2+^ (paired t-test, p < 0.0001, n = 11), indicative of an increase in channel activity, and when 1 µM LY354740 was added to the bath the variance decreased (paired t-test, p = 0.0233, n = 11), indicative of a decrease in channel activity (**Fig. 5B**). When the NMDG solution was applied, there was no significant change in variance, indicating that activating group II mGluRs had caused the closure of a Na^+^ conducting channel (paired t-test, p = 0.1002, n = 10).

In NALCN KO OFF UBCs, the effect of the group II mGluR agonist was absent (4.259 ± 5.204 pA, n = 7, paired t-test, p = 0.6564, **Fig. 5C**) and the amplitude of the NALCN current in NALCN KO OFF UBCs (-18.94 ± 6.999 pA, n = 7) was significantly less than in the wild type OFF UBCs (unpaired t-test, p = 0.0364, **Fig. 5D**) before significant reduction of current in NMDG (10.90 ± 3.184 pA, n = 7, paired t-test, p = 0.0141). The remaining inward current evoked by low Ca^2+^ in the NALCN KO could be due to incomplete removal of NALCN from the cells in this conditional knockout approach, or another current that is enhanced by the reduction of Ca^2+^.

NALCN can be modulated through G-proteins or through other intracellular pathways (Swayne *et al*., 2009; Monteil *et al*., 2024). To test whether NALCN inhibition by group II mGluRs is G-protein dependent, 1 mM GDP-β-S was added to the internal pipette solution to block the activation of intracellular G proteins. When NALCN was activated by the low calcium solution and 1 µM LY354740 was added to activate group II mGluRs, there was no significant reduction of the inward current in the presence of GDP-β-S (4.881 ± 2.206 pA, n = 11, paired t-test, p = 0.0514, **Fig. 5E**), followed by a significant reduction of the NALCN current remained when the solution was replaced with NMDG (43.36 ± 3.11 pA, n = 11, paired t-test, p < 0.0001). The amplitude of NALCN inhibition by LY354740 in WT OFF UBCs was significantly larger than the NALCN KO OFF UBCs (unpaired t-test, p = 0.0062) or when GDP-β-S was in the pipette solution (unpaired t-test, p = 0.0056). This indicates that group II mGluRs inhibit NALCN through the action of G proteins. Thus, group II mGluRs can inhibit NALCN in addition to activating GIRK. These mechanisms may allow glutamate to have a strong inhibitory influence on OFF UBCs to produce their long pauses in firing during and after high frequency synaptic stimulation.

### OFF UBCs have group III mGluRs that activate GIRK but do not inhibit NALCN

In addition to group II mGluR expression, RNA sequencing has shown group III mGluR expression in UBCs that, to our knowledge, has not been explored (Saunders *et al*., 2018; Kozareva *et al*., 2021; Jing *et al*., 2025). To test the physiological function of group III mGluRs in UBCs, we bath applied the agonist L-AP4 (Thomsen *et al*., 1992; Graham & Burgoyne, 1994; Thoreson & Ulphani, 1995). Application of 10 µM L-AP4 produced an outward current in the presence of 1 μM LY341495 (paired t-test, p = 0.0261, n = 8 cells, **Fig. 6A**). In current clamp, puff application of 300 µM L-AP4 hyperpolarized the cell and paused firing (**Fig. 6B**).

**Figure 6:**
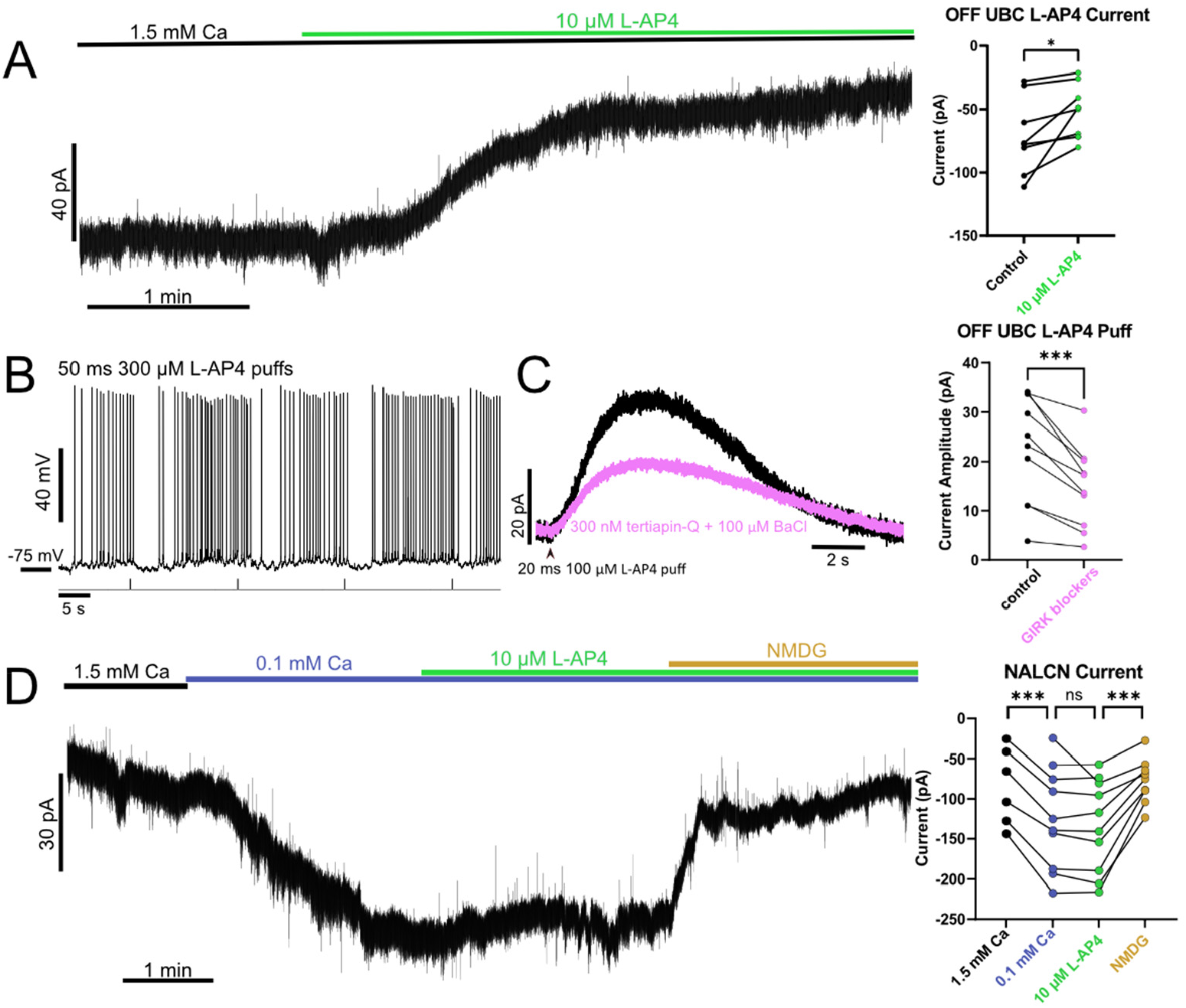
Group III mGluRs inhibit UBCs through activation of GIRK but not inhibition of NALCN. A) Left-voltage clamp recording of an OFF UBC. Bath application of 10 µM L-AP4 produced an outward current. Right-summary data. B) Brief puff application of 300 μM L-AP4 onto UBC dendritic brush hyperpolarizes the cell and pauses spontaneous firing in the presence of 1 μM LY341495 in current clamp recordings. C) Left-100 µM puff application of L-AP4 produced an outward current in voltage clamp that was partially blocked by GIRK antagonists (300 nM tertiapin-Q and 100 µM BaCl). Right-summary data. D) Left-Voltage clamp recording of an OFF UBC. Bath Ca^2+^ concentration was lowered from 1.5 mM to 0.1 mM to activate NALCN, then 10 µM L-AP4 was applied with no significant effect. Replacement of Na^+^ with NMDG confirmed the inward current was carried by Na^+^. Right-summary data.

To test if the hyperpolarizing effect of group III mGluRs was through the activation of GIRK, we puffed L-AP4 on OFF UBCs and applied GIRK blockers Tertiapin-Q and Ba^2+^ (**Fig. 6C**). Adding these GIRK blockers significantly decreased the amplitude of the L-AP4 evoked outward current (7.82 ± 1.51 pA, n = 10 cells, paired t-test, p = 0.0006, **Fig. 6C**). GIRK is difficult to block entirely with extracellular agents, so it is not clear whether the residual current was due to unblocked GIRK or another channel. Since we showed that group II mGluR activation inhibits NALCN along with GIRK, we ran similar experiments to test group III inhibition of NALCN. In the presence of GIRK blockers, an inward NALCN current was evoked by reducing the concentration of Ca^2+^ in the bath to 0.1 mM Ca^2+^ (-54.73 ± 5.231 pA, n = 6, paired t-test, p = 0.0001), then the group III mGluR agonist L-AP4 was applied **(Fig. 6D)**. L-AP4 application had no effect on the NALCN current (paired t-test, p = 0.2226, n = 10). Reduction of extracellular Na^+^ by replacement with NMDG confirmed that lowering Ca^2+^ had increased a Na^+^ leak current (61.87 ± 10.74 pA, n = 9, paired t-test, p = 0.0004, **Fig. 6D**). In sum, group III mGluRs do not inhibit NALCN, in contrast to group II mGluRs in the same cell type, suggesting distinct effector pathways for these GPCRs.

### GABA-B receptor activation in ON UBCs does not inhibit NALCN

Activation of GABA-B receptors has been reported to inhibit NALCN in dopaminergic cells in the substantia nigra and glycinergic cartwheel cells in the dorsal cochlear nucleus (Philippart & Khaliq, 2018; Ngodup *et al*., 2024). First we quantified the GABA-B current that has been previously reported in ON UBC (Kim *et al*., 2012; Guo *et al*., 2021; Huson & Regehr, 2025). The GABA-B agonist baclofen was bath applied in voltage clamp with a potassium gluconate based internal solution. Baclofen induced a small but consistent outward current (paired t-test, 5.869 ± 0.789 pA, p = 0.0001, n = 8, **Fig. 7A**). Next, to test whether GABA-B inhibits NALCN, extracellular Ca^2+^ was lowered to produce an NALCN current in the presence of GIRK blockers, then baclofen was applied to test whether it would reduce the inward current. NALCN was activated in low Ca^2+^ (-18.64 ± 3.839 pA, n = 11, paired t-test, p = 0.0007), but baclofen had no apparent effect on it (paired t-test, p = 0.2147, n = 11, **Fig. 7B**). The significant reduction in the NALCN current in the NMDG solution confirmed the inward current was carried by Na^+^ (15.78 ± 3.291 pA, n = 11, paired t-test, p = 0.0007). These data show that GABA-B receptors do not inhibit NALCN in ON UBCs, in contrast to group II mGluRs in OFF UBCs.

**Figure 7:**
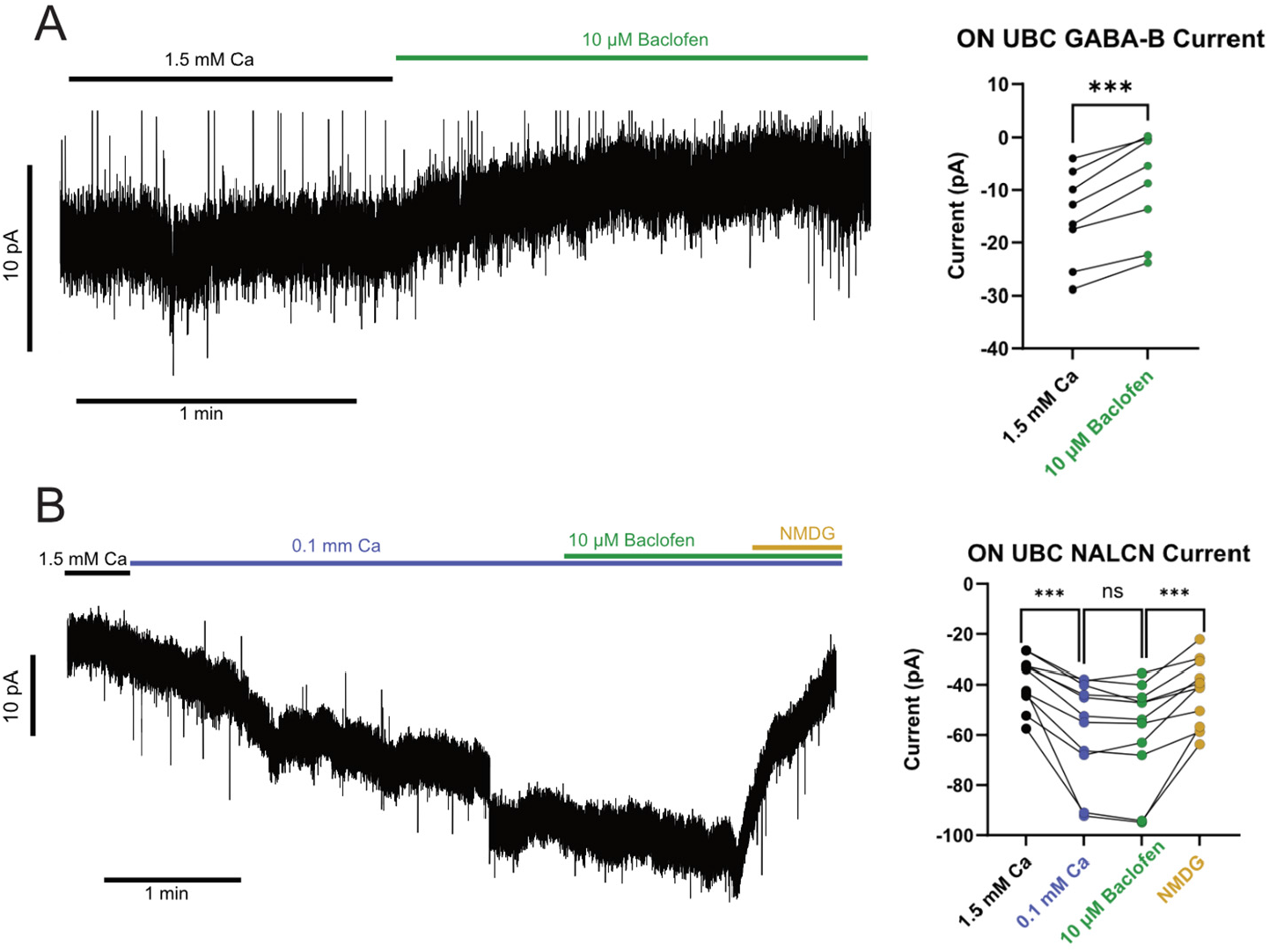
GABA-B activation does not inhibit NALCN in ON UBCs. A) Left-voltage clamp recording of GABA-B current in an ON UBC. 10 µM Baclofen was added to 1.5 mM Ca^2+^ ACSF and a modest inhibitory outward current was observed. Right-summary data. B) Left-voltage clamp recording of activation of NALCN current in an ON UBC. Solution was lowered to 0.1 mM Ca^2+^ and a depolarizing inward current was observed. When 10 µM baclofen was added to the bath it did not block the inward NALCN current. Applying the NMDG solution reduced the Na^+^ and returned the cell to baseline. Right-summary data.

## Discussion

### A novel mechanism of glutamatergic synaptic inhibition

Although glutamate typically excites neurons, the action of glutamate on metabotropic receptors depends on the intracellular pathways that are initiated. In retinal bipolar cells, mGluR6 has an inhibitory effect by activating GIRK (Nawy & Jahr, 1990; Tian & Kammermeier, 2006; Rai *et al*., 2021). Group II mGluRs in various neuron types also activate GIRK, but whether they can influence NALCN currents had not been tested. Here we show that in OFF UBCs group II mGluRs inhibit a tonic NALCN current, in addition to activating GIRK. These actions together produce the large hyperpolarizing current in OFF UBCs. This mechanism is likely present in other cells that express group II mGluRs and NALCN together, such as neurons in the olfactory bulb, hippocampus, thalamus, parietal and somatosensory cortex, and other cerebellar neurons (Ohishi *et al*., 1998).

To our knowledge, this is a newly discovered mechanism of glutamatergic synaptic inhibition. Synaptic inhibition is most typically due to the opening of a channel that conducts anions into the cell (e.g. Cl^-^ channels) or cations out of the cell (e.g. K^+^ channels). Shunting inhibition is also common, in which channel opening reduces the input resistance of the cell, making excitatory currents less effective in changing the membrane potential. In contrast, this mechanism of synaptic inhibition in which glutamate closes an open channel that conducts an inward current, would hyperpolarize the cell while simultaneously increasing its input resistance. The increase in resistance would enhance the cells response to synaptic potentials. Thus, this mechanism is expected to reduce the firing of a neuron, while at the same time, enhancing its sensitivity to subsequent postsynaptic potentials by increasing their amplitude and slowing their decay. Inhibition of NALCN through this mechanism could modulate how a cell transforms signals by changing its firing phase, duration, or properties of integration. This mechanism could broaden the computational processing ability of UBCs and influence the transformation of vestibular signals that underlie balance and eye movements. Other Gi/o-coupled receptors present in many neuron types could have similar inhibitory actions by blocking additional tonically active inward currents, such as persistent sodium currents, HCN currents, TRP channel currents, or Ca^2+^ channel currents (Han *et al*., 2006).

### NALCN as a regulator of rhythmic firing

The tonic inward Na^+^ current has been implicated in the rhythmic firing of several neuron types (Lu *et al*., 2007; Shi *et al*., 2016; Um *et al*., 2021; Wang *et al*., 2023; Cobb-Lewis *et al*., 2023). We show that OFF UBCs have a larger tonic inward NALCN current than ON UBCs, and this has a depolarizing influence on their resting membrane potential. OFF UBCs have a more regular spontaneous firing pattern than ON UBCs (Russo *et al*., 2007; Diana *et al*., 2007; Kim *et al*., 2012; Balmer *et al*., 2021). We found that OFF UBCs lacking NALCN have a more irregular and less rhythmic firing pattern, which suggests that this inward leak current is a mechanism to enforce consistent interspike intervals. The difference in the level of NALCN current in ON and OFF UBCs may determine their firing regularity. These data add to the growing body of evidence that NALCN is important for rhythmic firing (Lu *et al*., 2007; Lutas *et al*., 2016; Um *et al*., 2021; Wang *et al*., 2023; Cobb-Lewis *et al*., 2023) and extend it to suggest that it may regulate the firing pattern as well.

Firing variability may prime neurons for different roles in sensory processing. Like ON and OFF UBCs, vestibular afferents can be classified by their regular and irregular firing patterns (Goldberg & Fernández, 1977; Lasker *et al*., 2008). Irregular vestibular afferents respond quicker to head movement stimuli than regular vestibular afferents and have a larger response to electrical stimulation than regular afferents (Goldberg *et al*., 1984; Goldberg, 2000). Meanwhile, regular firing afferents are better suited for slower movements having lower thresholds, less adaptation, and longer duration responses (Sadeghi *et al*., 2007; Eatock *et al*., 2008). Various ion channels are responsible for promoting regular and irregular firing in vestibular afferents (Eatock *et al*., 2008). NALCN could be an additional conductance that controls the regularity of various neurons including vestibular afferents to tune them to process different types of sensory input.

In the vestibular cerebellum, OFF UBCs receive secondary vestibular signals from the vestibular nucleus, whereas ON UBCs receive both secondary and primary afferent input from the vestibular ganglion (Balmer & Trussell, 2021).

Primary afferents likely fire faster than secondary afferents, so we hypothesize that the irregular firing of ON UBCs functions to allow rapid processing of these sensory signals. This would be an effective way for each ON UBC to amplify primary vestibular signals from a single vestibular organ to an ensemble of hundreds of granule cells via its branching axons and mossy fiber-like terminals.

### Group III mGluRs and GABA-B receptors do not inhibit NALCN in UBCs

We show for the first time that UBCs are hyperpolarized by group III mGluRs. We tested whether these receptors act through inhibition of NALCN, but unlike group II mGluRs (mGluR2/3), we find that these receptors produce inhibition through GIRK but not NALCN. Since group III receptors are commonly present on axons (Shigemoto *et al*., 1997), we confirmed that they are present on UBC dendrites by focal agonist puffs. GABA-B receptors also had no apparent effect on NALCN in UBCs, although GABA-B has been shown to inhibit NALCN in dopaminergic neurons in the substantia nigra and cartwheel cells in the dorsal cochlear nucleus (Philippart & Khaliq, 2018; Ngodup *et al*., 2024). Group II mGluRs may inhibit NALCN because they are physically closer to NALCN than are group III mGluRs and GABA-B receptors. The subcellular location of NALCN has been shown to be important in substantia nigra neurons, in which NALCN channels located at proximal dendritic sites are crucial for burst firing (Hahn *et al*., 2023). Further biochemical studies to determine the proximity of NALCN to these GPCRs may address how the same G-proteins target different ion channels in the same cell.

### G-protein dependent mechanisms of NALCN modulation

NALCN channels were inhibited by a group II mGluR agonist, and this effect was occluded by the presence of intracellular GDP-β-S, indicating that group II mGluR inhibition of NALCN is G-protein dependent. NALCN has been shown to be activated and inhibited by G-protein independent pathways as well. For example, NALCN orthologs in C. elegans can be activated by Gq-rho (Topalidou *et al*., 2017). In pancreatic β-cells, muscarinic acetylcholine receptors form a complex with NALCN and gate them in a G protein-independent manner. In ventral tegmental and hippocampal neurons, peptide neurotransmitters substance P and neurotensin activate NALCN through a Src kinase pathway (Lu *et al*., 2009).

NALCN is present in many non-neuronal cells and nearly all neurons and multiple pathways of positive and negative modulation may have evolved to allow various receptors to have effects on cellular excitability with different time scales from milliseconds (neurotransmitters) to seconds (neuromodulators) and hours (circadian rhythms) (Gonzalez *et al*., 2023).

## Conclusion

This work reports a previously undescribed mechanism through which glutamate can inhibit neurons. This form of inhibition may differ from classical inhibition, because closing NALCN channels could increase membrane resistance and enhance subsequent synaptic potentials, which is opposite to the shunting effect caused by opening Cl^-^ or K^+^ channels. We also show that NALCN enforces regular firing in OFF UBCs, and we hypothesize that the relatively lower NALCN current in ON UBCs may contribute to their irregular firing. This mGluR-NALCN mechanism of synaptic inhibition may be present in neurons across the nervous system and may be relevant to disorders of excitability including epilepsy, cerebellar ataxia, and tinnitus (Aoyagi *et al*., 2015; Nguyen *et al*., 2023; Cabrita Pinto *et al*., 2025).

## Author Contributions

C.T.C., K.E.W., and T.S.B. designed and analyzed the experiments, and contributed to writing the manuscript, C.T.C., K.E.W. executed experiments, T.S.B. supervised. All authors have read and approved the final version of this manuscript and agree to be accountable for all aspects of the work in ensuring that questions related to the accuracy or integrity of any part of the work are appropriately investigated and resolved. All persons designated as authors qualify for authorship, and all those who qualify for authorship are listed.

## Data Availability

The data that support the findings of this study are available upon reasonable request.

